# A fluorogenic nanobody array tag for prolonged single molecule imaging in live cells

**DOI:** 10.1101/111690

**Authors:** Rajarshi P. Ghosh, Will E. Draper, J. Matthew Franklin, Quanming Shi, Jan T. Liphardt

## Abstract

Prolonged single molecule imaging in live cells requires labels that do not aggregate, have high contrast, and are photo-stable. To address these requirements, we have generated arrays of modular protein domains that function as fluorophore recruitment platforms. ArrayG, a linear repeat of GFP-nanobodies, recruits free monomeric wild-type GFP, which brightens ~15-fold upon binding the array. The fluorogenic ArrayG tag effectively eliminates background fluorescence from free binders, a major impediment to high-throughput acquisition of long trajectories in recruitment based imaging strategies. The photo-stability of ArrayG and consistently low background made it possible to continuously track single integrins for as long as 105 seconds (2100 frames). Prolonged tracking of both kinesin and integrin revealed repeated state-switching events, a measurement capability that is crucial to a mechanistic understanding of complex cellular processes. We also report an orthogonal array tag, based on a DHFR-nanobody, for prolonged dual color imaging of single molecules.

## Introduction

Single molecule experiments in live cells can reveal the operating principles of cellular machineries^1,2^, but frequently present challenges including fluorophore delivery, conjugation, and intrinsic cellular fluorescence.^3,4^ Fluorescent proteins (FP) expressed as direct fusions to cellular proteins address delivery and specificity, but do not emit enough photons for prolonged single-molecule tracking. Trajectory duration sets a fundamental limit on the description of a biological system, most acutely in situations where molecules exhibit distinct behaviors as they traverse multiple microenvironments and interact with distinct binding partners.

One way to overcome the limitations of FPs as single molecule probes is to fuse multiple copies of genetically encoded FPs directly to a target^5^ or though recruitment to arrays. For example, the behaviors of single RNA molecules and DNA loci have been monitored using FP fusions to nucleic acid binding proteins that are recruited to cognate sequence arrays.^6-8^ The recently developed ‘SunTag’ protein label is an array of short epitopes that binds to single-chain GCN4 antibodies (scFv-GCN4).^9^ One limiting feature of these systems is the difficulty of dialing in relative expression levels of the fluorescence binders and their target arrays, such that array occupancy is maximized and yet background signal due to free binders is minimized.

Lately there has been a surge in the development of single-chain, high affinity protein-binding domains such as Nanobodies,^10^ DARPins,^11^ Monobodies,^12^ and Affibodies.^13^ Starting with an initial candidate set of six array/ binder pairs, five of which are nanobody based, we have selectively developed two genetically encoded tags for prolonged imaging of single molecules in living cells. These arrays consist of repeated units of camelid nanobodies.^14^ We show that array concatenation retains important functionalities of the constituent domains, and therefore can act as more than simple recruitment platforms. For example, one of these nanobody arrays enhances wild-type GFP (wtGFP) fluorescence upon binding, resulting in a consistently high signal-to-noise ratio due to minimal background fluorescence of the free binder. This enabled high throughput acquisition of single molecule images. Development of a second orthogonal nanobody based array system allowed simultaneous multicolor imaging. We show that these arrays make it possible to follow the behavior of single kinesin and integrin molecules in living cells for long times, with no visible perturbation of each molecule's native function. Utilizing the long-term photo-stability of these arrays we efficiently captured the multi-state dynamics of both kinesin and integrin molecules, demonstrating the utility of these tools for dissecting complex mechanistic questions inside live cells.

## Results

### Motivation for a ‘dynamic’ array strategy

The simplest strategy for amplifying fluorescence from a single molecule is to fuse multiple copies of a fluorescent protein in a linear sequence. We tested whether fusing 20 copies of super-folder GFP (sfGFPs) resulted in a stoichiometric increase in its fluorescence (sfGFP_20x_, **Figure 1a-c**). When expressed in HeLa cells and imaged using HiLo^15^ TIRF microscopy, the average intensity of sfGFP_20x_ spots corresponded to approximately five purified eGFPs (**Figure 1b, Methods**). Previous research has found little evidence for GFP self-quenching^16,17^ or hypo-chromicity,^18^ suggesting that the weak fluorescence of sfGFP_20x_ reflected impaired folding or maturation.

**Figure 1.**
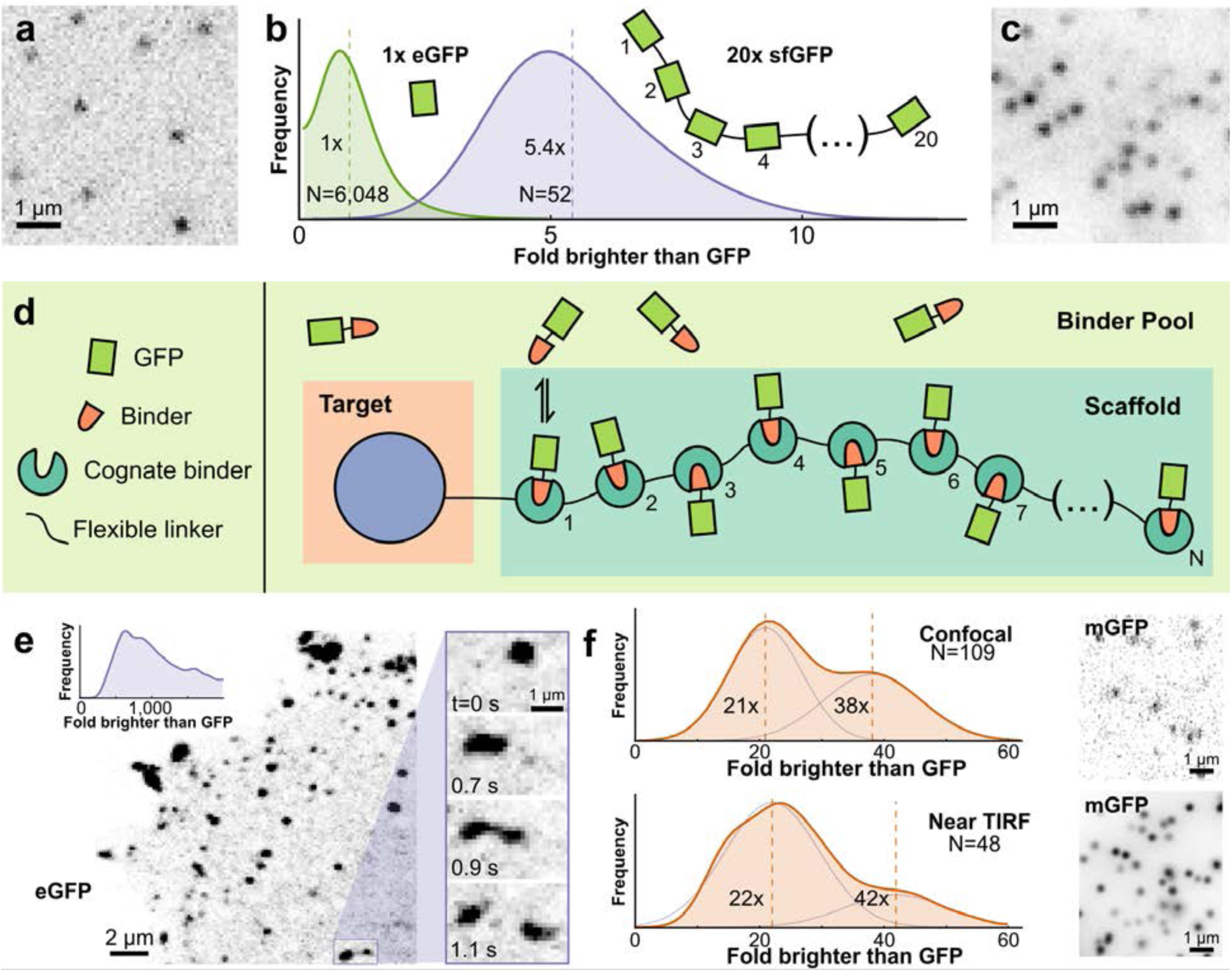
Approach for prolonged imaging and abolishing aggregation propensity. (**a**) Representative image of single purified eGFP on glass imaged using TIRF microscopy. (**b**) Distribution of intensities of 20x-sfGFP (super-folder GFP, blue curve) expressed in HeLa cells presented as fold-brighter than mean eGFP intensity (green curve). (**c**) Representative image of membrane-proximal 20x-sfGFP molecules imaged using identical settings as in (a). (**d**) Schematic of the prolonged imaging *in cellulo* approach. (e) Representative laser-scanning confocal image of a cell co-expressing kinesin-1(KIF560)-ArrayG (24x GBP1 GFP-nanobody array) along with eGFP as the binder. Right inset: Zoom-in showing droplet-like temporal dynamics of kinesin aggregates. Top left inset: Distribution of intensities of spots in terms of fold brighter than eGFP, showing extensive aggregation. The x axis has been truncated at 2,000 eGFPs to emphasize the large size of even the smallest aggregates. (**f-g**) The use of monomeric A206K GFP (mGFP) abolishes aggregation. KIF560-ArrayG intensity distributions represented as fold-brighter than eGFP, show peaks corresponding to KIF560-ArrayG monomer and dimer for confocal (top) and Hilo-TIRF (bottom) (**f**), with representative images (**g**). Intensity distributions were fit to a two-Gaussian mixture model to estimate the local maxima for the monomeric and dimeric peaks. Vertical lines in intensity distributions represent the mean value.

Although these problems are in principle addressable via protein engineering, we pursued a more general solution based on dynamic recruitment of fluorescent proteins or protein fusions to a repeat array (**Figure 1d**). Previous work has demonstrated the utility of this approach using peptide epitope arrays^9^ and we decided to extend this approach to protein domains so that we could leverage binding-dependent emergent properties such as fluorogenicity. Simultaneous usage of multiple arrays with different cognate binders would also allow multicolor imaging.

### Preliminary scan of possible array/binder pairs

We surveyed the literature for reasonably small, high affinity protein interaction pairs that were unlikely to interfere with normal cellular processes. We identified six potentially suitable pairs ranging from short peptide epitopes to larger protein domains (**Supplementary Table 1**). Of those selected, five were nanobody-antigen pairs. The small size and monomeric nature of nanobodies makes them particularly suitable for generating tandem repeat arrays for protein tagging.^10^ The *in cellulo* recruitment potential of each interaction pair was assessed using a nuclear compartmentalization assay. This assay tested the ability of a histone H2B fusion of a protein domain, expressed either as a single copy or as a 24X repeat array, to sequester its cognate binder in the nucleus (**Methods, Supplementary Figure S1, Supplementary Table 1**). From these assays we chose the best performing interaction pairs: GBP1-GFP and NB113-DHFR. As our first array, we chose the GBP1-GFP binder pair, which provided a potentially attractive candidate for background fluorescence suppression, based on the previously reported role of GBP1 in enhancing wtGFP fluorescence.^19^

### Eliminating aggregation propensity

We began our evaluation of the GBP1 nanobody by fusing a 24x repeat (GBP1_24x_) to KIF560, a (-)-end directed kinesin-1 mutant lacking the cargo-binding domain.^5,20,21^ When co-expressed with eGFP, we observed numerous bright fluorescent spots exhibiting complex dynamic behavior including directed motion (**Supplementary Movie S1**). We used calibrated confocal microscopy to analyze the spot intensity distribution, which revealed that all detectable spots were large aggregates (**Figure 1e, Supplementary Figure S2**). The smallest detectable KIF560- GBP1_24x_ spots corresponded to ~250 eGFPs, with a typical spot containing ~1000 eGFPs (**Figure 1e inset**).

Since eGFP has a weak dimerization propensity (K_d_ = 110 μM)^22^, we tested whether the strictly monomeric A206K^22^ mutant of eGFP (‘mGFP’) would eliminate aggregation. Remarkably, replacing eGFP with mGFP collapsed the formerly broad intensity histograms onto sharp two-peaked distributions with maxima corresponding to 21 and 38 GFPs (confocal microscopy) or 22 and 42 GFPs (HiLo-TIRFM) (**Figure 1f-g**). The two peaked distribution likely reflects dimers of kinesin with either one or two GBP1_24x_ labels per-molecule.^23^

We further tested the GBP1_24x_–mGFP combination by examining the diffusion of free GBP1_24x_ arrays. We tracked the arrays using HiLo-TIRFM at 100 Hz (**Supplementary Figure S3, Supplementary Movie S2**). The free GBP1_24x_ arrays had an apparent diffusion coefficient of 0.30 μm^2^/s, showing no evidence of aggregation. As expected for inert tracers of this size (~1 MDa) such as Dextran, these exhibited sub-diffusive behavior (**Supplementary Figure S3**).^24^

Although the GBP1_24x_-mGFP pair allowed single molecule tracking in living cells, the variation in the GBP1_24x_ to mGFP expression ratio greatly reduced experimental throughput. One approach to this problem is to sequester the freely diffusing FP population into the nucleus using an NLS^9^. However, this might increase the effective size of the tag due to binding by importins. An alternative approach is to sort for cells of low fluorescence, but this is not amenable to high-throughput testing of target fusions and sensitive cell lines. Further, the low concentration of free binders may result in under-occupied arrays. The ideal strategy for background fluorescence suppression is to use a fluorogenic system, where the free fluorophore is essentially dark and becomes highly fluorescent upon binding to the target array (**Figure 2a**).

**Figure 2.**
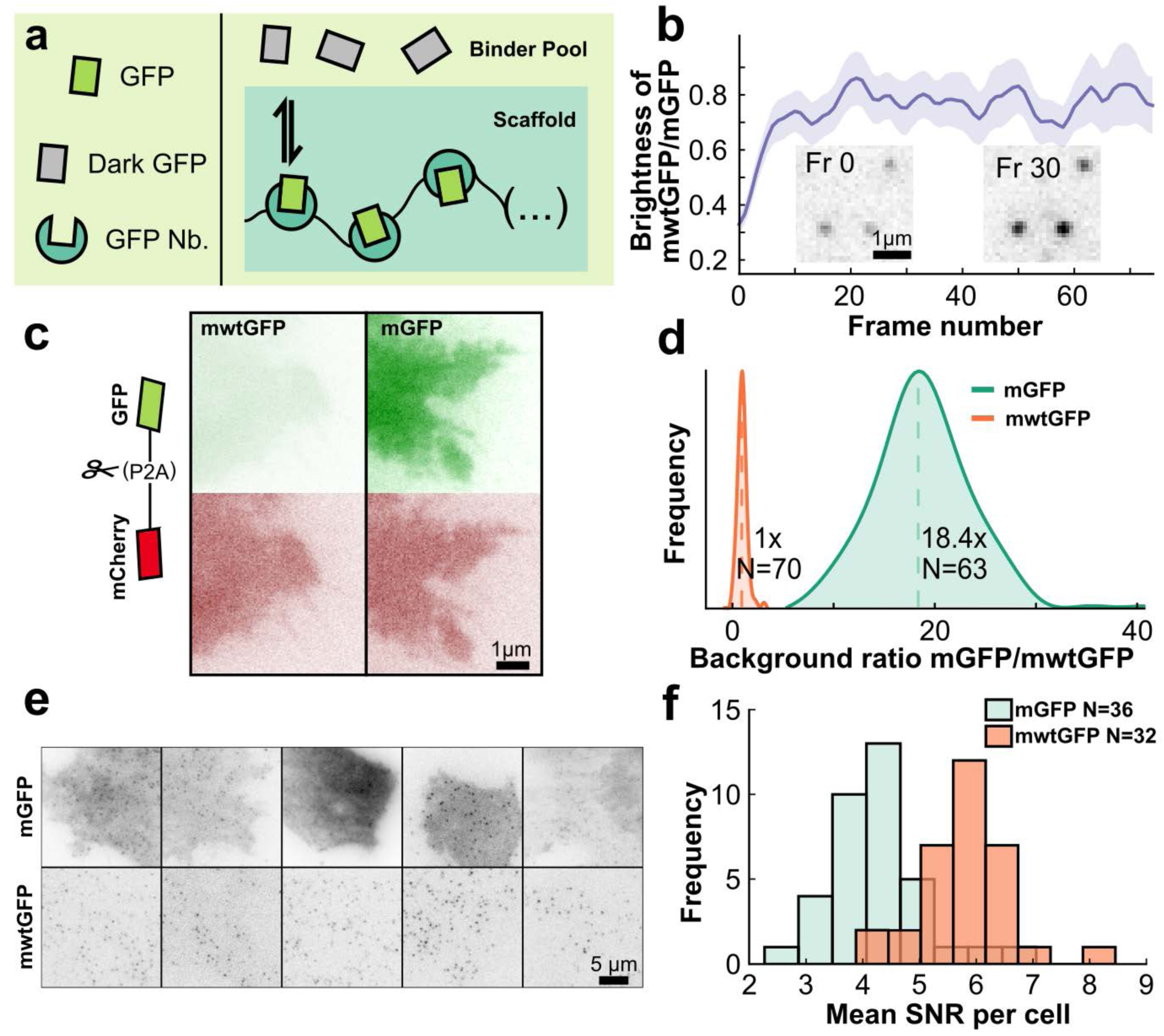
Signal enhancement through background reduction by GBP1 binding-induced fluorogenicity. (**a**) Schematic of binding-induced fluorescence enhancement. (**b**) Time dependence mwtGFP *(N* = 5564 traces) to mGFP *(N* = 1862 traces) spot brightness ratio. Inset: TIRF images of mwtGFP spots at frame 0 and frame 30 (at 20Hz image frequency), showing significant brightening. (**c)** TIRF images of HeLa cell lines stably expressing self-cleavable constructs for ratiometric normalization of GFP fluorescence: mCherry-p2a-mwtGFP (left) or mCherry-p2a-mGFP (right). (**d**) Distribution of mGFP intensity *(N =* 63 cells) normalized by mean mwtGFP intensity *(N =* 70 cells) measured using TIRF. Vertical dashed lines represent mean value. (**e**) TIRF images of U2OS cells stably expressing integrin β1-ArrayG_16x_ and either mGFP (top) or mwtGFP (bottom) binder. (**f**) Histogram of the mean particle signal-to-noise per cell for integrin β1-ArrayG_16x_ co-expressing either mGFP or mwtGFP.

### Binding induced fluorescence enhancement for background suppression

GBP1 has been previously shown to enhance the fluorescence of wtGFP upon binding.^19^ We hypothesized that using wtGFP instead of mGFP would greatly reduce background fluorescence resulting from free binders. To make the wtGFP stringently monomeric we introduced the monomeric A206K mutation (mwtGFP).

To measure the extent of the fluorogenic effect in the context of the GBP1 array, we co-expressed KIF560-GBP1_24x_ fusions in cells with either mwtGFP or mGFP. GBP1_24x_ was ~75% as bright when occupied with mwtGFP than when occupied with mGFP (**Figure 2b**). An unexpected finding was that mwtGFP-bound GBP1_24x_ brightened upon illumination. This photo-maturation occurred even at relatively low laser powers (< 0.01 kW cm^2^) and was complete within ~200–300 ms.

To quantitatively compare fluorescence between cells expressing mwtGFP and mGFP, we generated HeLa cell lines expressing either mwtGFP or mGFP fused to mCherry via a P2A^25^ self-cleavable peptide bridge (**Figure 2c**). This system allowed us to measure the relative per-molar GFP fluorescence by normalizing to mCherry fluorescence. Flow cytometric analysis of the normalized basal fluorescence showed ~30 fold less fluorescence for free mwtGFP compared to mGFP (**Supplementary Figure S4**). In contrast to FACS, HiLo-TIRFM imaging revealed an 18-fold weaker fluorescence for mwtGFP (**Figure 2d**). This difference in relative fluorescence levels is most likely due to the photo-maturation effect of mwtGFP upon continuous illumination.

We hypothesized that by using mwtGFP, which has near equivalent brightness to mGFP (~75%) and substantially low (18 fold) background, we would obviate the need for background fluorescence optimization, thus increasing the frequency of finding cells with high signal-to-noise ratio (SNR). To test this possibility, we generated U2OS cell lines co-expressing the integral membrane protein, integrin β1, fused to a GBP1_16x_ array, under cumate inducible control and either mwtGFP or mGFP under doxycycline inducible control. We chose integrin β1, since this would spatially confine the GBP1 scaffold to a single cellular compartment, facilitating quantitative comparison.

After inducing mGFP or mwtGFP, nearly every cell co-expressing integrin β1-GBP1_16x_ and mwtGFP produced detectable single molecules exhibiting high signal over background (**Figure 2e**). However, cells co-expressing β1-GBP1_16x_ and mGFP exhibited high variability in background signal, which in some cells obscured the detection of single molecules altogether (**Figure 2e**). To quantify this difference, the mean signal-to-noise ratio (SNR) of localized diffraction limited spots on a per-cell basis was measured for integrin β1-GBP1_16x_ with either mGFP (*N* = 36 cells) or mwtGFP (*N* = 32 cells) under identical imaging conditions. The use of mwtGFP resulted in a significant increase in SNR from a mean of 4.3 for mGFP to 5.8 for mwtGFP (**Figure 2f**). This shift in mean SNR is a considerable underestimation of the SNR enhancement achieved through the use of mwtGFP, since many of the mGFP cells had too high a background for detection of single-molecules and therefore were not included in the comparative SNR analysis.

For the purpose of prolonged single molecule tracking inside cells, the benefits of a near-zero background far outweigh the 25% reduction of brightness caused by replacing mGFP with mwtGFP. The GBP1_24x_ / mGFP pair is a viable option where the very highest localization precisions are required, at the expense of severely reducing the number of cells with single molecules that have a high SNR. For ease of discussion, we refer to the GBP1_24x_/mwtGFP combination simply as ‘ArrayG_24x_’.

### Speed variations and hopping behaviors of kinesin

Having developed an array with no detectable aggregation propensity and consistently high SNR, we repeated the imaging of KIF560 with the optimized ArrayG_24x_. Single particle tracking (SPT) of KIF560-ArrayG_24x_ using HiLo-TIRFM (20 Hz) yielded hundreds of directionally processive trajectories per cell, spanning up to several μm in length (**Figure 3a-b**). The speed of KIF560-ArrayG_24x_ motors at 37 °C was sharply distributed about a mean of 1.48 μm/s (**Figure 3b-c**, **Supplementary Movie S3**). Although the large size of ArrayG_24x_ could in principle affect Kinesin’s *in cellulo* behavior, we did not detect any negative repercussions, as the labeled kinesins recapitulated previously reported average speeds and run-lengths.^5,9^

**Figure 3.**
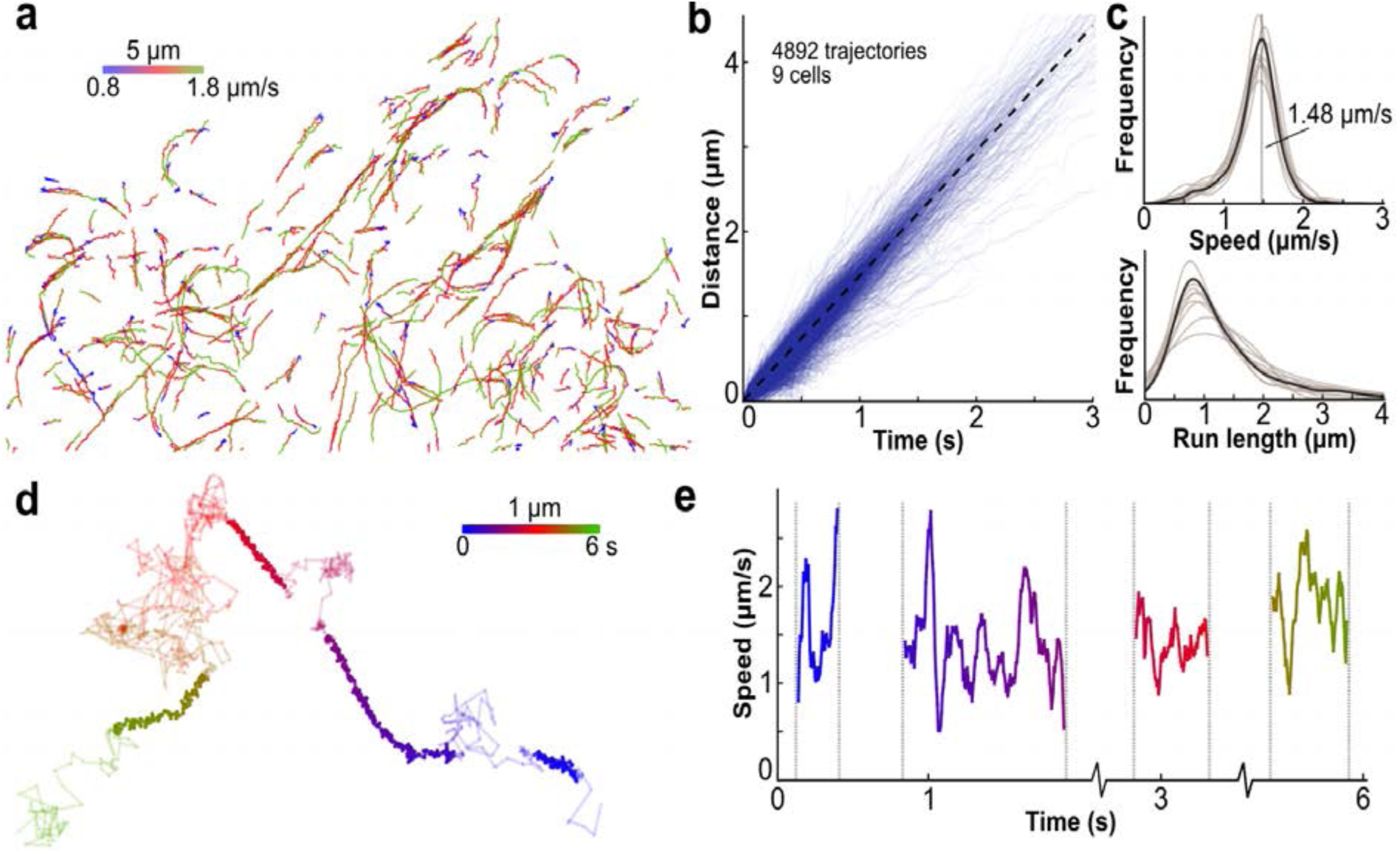
Analysis of Kinesin-ArrayG trajectories demonstrates proper functionality and reveals fast state-switching. **(a)** Trajectories of single KIF560-ArrayG motors labeled with mwtGFP colored by speed. **(b)** Distance versus time for 4892 individual trajectories from 9 cells. Dashed line represents the mean speed. **(c)** Distribution of trajectory speeds (top) and run lengths (bottom). Individual cell distributions are shown in transparent, with the mean as solid. Vertical dashed line indicates the mean (1.5 μm/s). **(d)** A high frame-rate trajectory of KIF560-ArrayG (180Hz) colored by time. Linearly persistent trajectory segments are colored bold. **(e)** Instantaneous speed of the linearly persistent segments from the trajectory shown in **d**.

By increasing the laser illumination 25-fold to 0.2 kW/cm^2^, single molecules of KIF560-ArrayG_24x_ were visible at 180 Hz. At such high frequencies, even short trajectories consist of hundreds of data points, allowing single trajectories to be analyzed in detail. In single molecule trajectories of KIF560- ArrayG_24x_, we observed multiple cycles of microtubule binding, directed motion along the microtubule, unbinding, and free diffusion. Several trajectories displayed multiple (up to 4) binding/run/unbinding/diffusion cycles over ~800 frames (**Figure 3d**, **Supplementary Movie S4**). During the periods of directed motion, the speed varied dramatically from less than 1 μm/s to nearly 3 μm/s, with an average speed around 1.5 μm/s, in accordance with the lower time resolution data (**Figure 3e**). To our knowledge, such transient variations in speed only detectable at very high time resolution, has not been previously reported using genetically-encoded tags. Two reports which have tracked quantum dot conjugated kinesins in live cells at relatively lower time resolution (10 to 30 Hz) differ in their fundamental findings regarding velocity and run lengths.^26, 27^

### Successive state transitions in integrin β1 trajectories revealed by prolonged imaging

To further explore the utility of the ArrayG tag, we studied the dynamics of integrin β1, a heterodimeric transmembrane protein. Previous single molecule experiments have suggested that integrins transition between diffusive inactive conformations and immobile active conformations, driven by binding to ECM ligands or through cytoplasmic interactions with focal adhesion components.^28^ Low time-resolution TIRFM imaging of synthetic dye labeled integrins (5Hz)^28^ and unconventional thermal imaging using gold nanoparticle labeled integrins^29^ have revealed state-switching. We reasoned that the ArrayG tag owing to its high photo-stability and biocompatibility would provide a convenient tool for long term imaging of integrins.

To assess the biological function of ArrayG-tagged integrin β1, we asked if the tagged integrin would restore cellular spreading of a β1 knockout cell line. The pan-knockout mouse embryonic fibroblast cell line pKOαV β1^-/-^ has reduced spreading area on fibronectin and this defect can be rescued by re-introduction of wildtype β1.^30^ We used the PiggyBac transposable vector system to stably express integrin β1-ArrayG under cumate control and mGFP under tetracycline control (**Supplementary Movie S5**). Preliminary comparison of β1-ArrayG_24x_ scaffolds to β1-GFP_2x_ fusions showed a lack of robust focal adhesion localization. To minimize interference with dynamics and localization, we fused integrin β1 to a shorter version of ArrayG (β1-ArrayG_16x_).

Expression of β1-ArrayG_16x_ in the β1^-/-^ cell line resulted in bright membrane-localized assemblies with the characteristic shape and location of focal adhesions (**Figure 4a**), indicating formation of the αβ heterodimer and proper localization. To assess whether β1-ArrayG_16x_ reverses deficient cell spreading in pKOαVβ1^-/-^ cells, we measured the average cell area over time for (1) pKOαV β1^-/-^ (negative control), (2) pKOαV β1^-/-^ + β1-P2A-eGFP (positive control) (**Supplementary Figure S5**) and (3) pKOαVβ1^-/-^ + β1- ArrayG_16x_ + mGFP (**Figure 4b**). The β1-P2A-eGFP construct self-cleaves to leave native integrin β1 and a diffusible eGFP as a marker for integrin β1 expression. Cells expressing either β1-ArrayG_16x_ or β1-P2A-eGFP had a large mean spreading area after 5 hours (1100 ± 470 μm^2^ and 1100 ± 390 μm^2^, mean **±** s.d.) whereas cells lacking β1 spread significantly less, as expected^30^ (560 ± 240 μm^2^, **Figure 4b, 4c, Supplementary Figure S5**), demonstrating that β1-ArrayG_16x_ can restore cell-spreading to wild-type levels.

**Figure 4.**
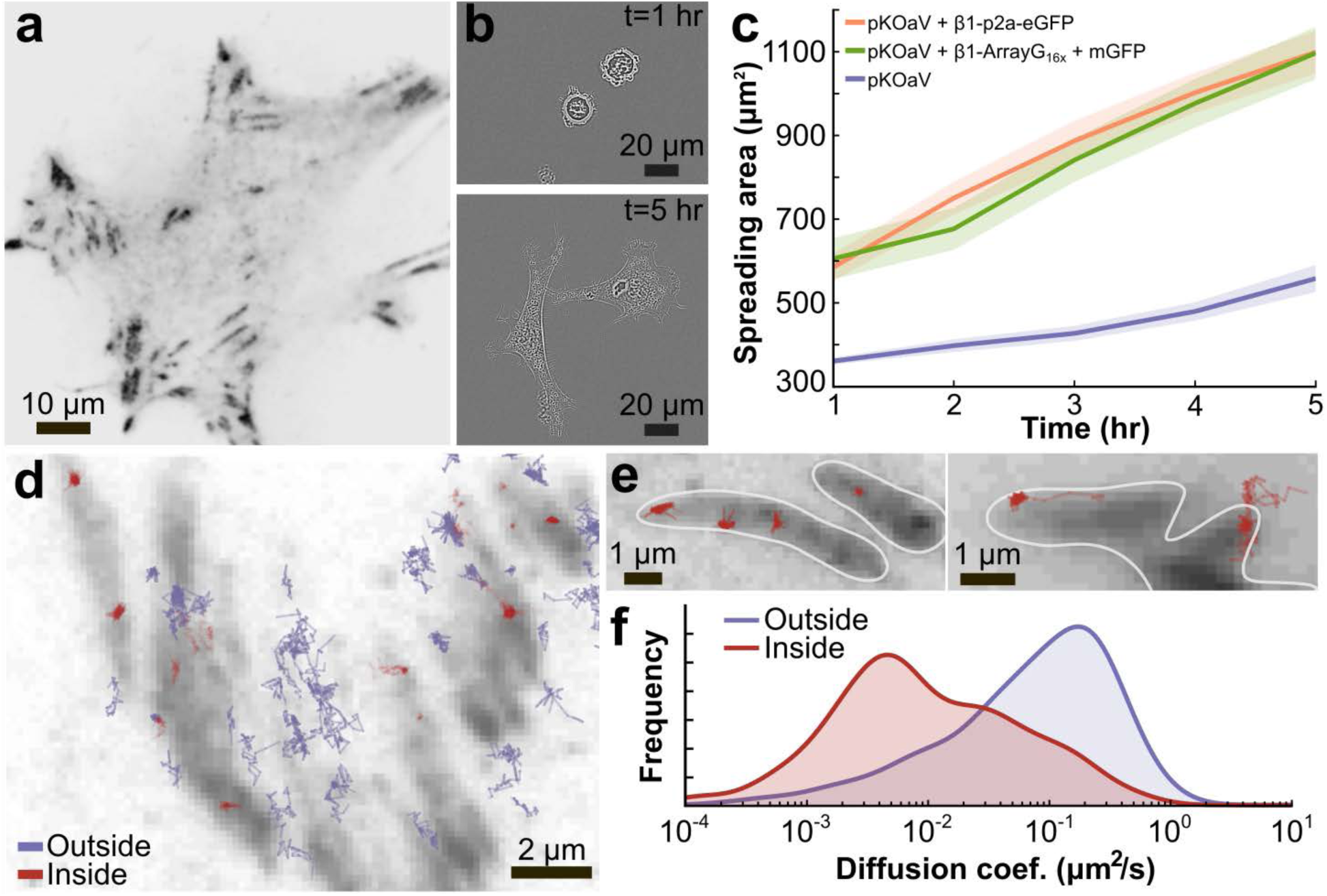
Evaluation of integrin β1-ArrayG_16x_ functionality. (**a**) Representative TIRF image of pKOαV MEF cells after induction of integrin β1-ArrayG_16x_ and mGFP. (**b**) Brightfield images of pKOaV MEFs overexpressing β1-ArrayG_16x_ + mGFP at 1 and 5 hours post seeding on fibronectin. (**c**) Mean spreading area versus time for *pKOαV* (53 cells), *pKOαV+β1-(p2a)-eGFP* (59 cells), and *PkoαV+β1- ArrayG*_16x_*+mGFP* (57 cells). Shaded area represents standard error of the mean. (**d**) β1-ArrayG_16x_ trajectories overlayed with time-averaged vinculin-mCherry in grey. Minimum trajectory length is 1 second. (**e**) Examples of immobilized trajectories inside focal adhesions (left) and trajectories transitioning between diffusive and immobilized states at focal adhesion boundaries (right). Focal adhesion boundaries highlighted in white. (**f**) Diffusion coefficient distribution of integrin β1-ArrayG_16x_ inside and outside focal adhesions. *N =* 5 cells, 1852 trajectory segments.

For validating functionality at the single molecule level, we first investigated whether β1-ArrayG_16x_ exhibited increased immobilization within cellular adhesions, as previously reported.^28^ In adhesions marked by vinculin-mCherry, we observed many strongly immobilized β1-ArrayG_16x_ molecules (**Figure 4d**). Further, we were able to directly capture the processes of lateral diffusion into adhesions and subsequent immobilization (**Figure 4e, Supplementary Movie S6**). Quantification of diffusivity revealed enhanced immobilization of integrin β1-ArrayG_16x_ molecules localized inside adhesions (**Figure 4f**). Despite the large size, the β1-ArrayG_16x_ traversed across adhesion boundaries, which is consistent with the suggested ‘archipelago’ like architecture of focal adhesions.^31^

Previous work has found that integrin clustering is required for cell spreading and traction force loading, suggesting integrin function is dependent on its local density.^32,33^ To assess whether a β1- ArrayG_16x_ is capable of processing external signals in a native background, we compared its dynamics to a simple β1-eGFP_2x_ fusion in pKOαVβ1 cells that express endogenous integrin β1. The diffusion coefficients had a bimodal distribution representing a freely diffusing and an immobilized population, in accordance with previous measurements^28^ (trajectories in **Figure 5a** and **Supplementary Figure S6**, distributions in **Figure 5b**). Mobile β1-eGFP_2x_ molecules had a slightly (~1.9 fold) larger diffusion coefficient compared to β1-ArrayG_16x_, but the overall shape and peaks of the two distributions were nearly identical.

**Figure 5.**
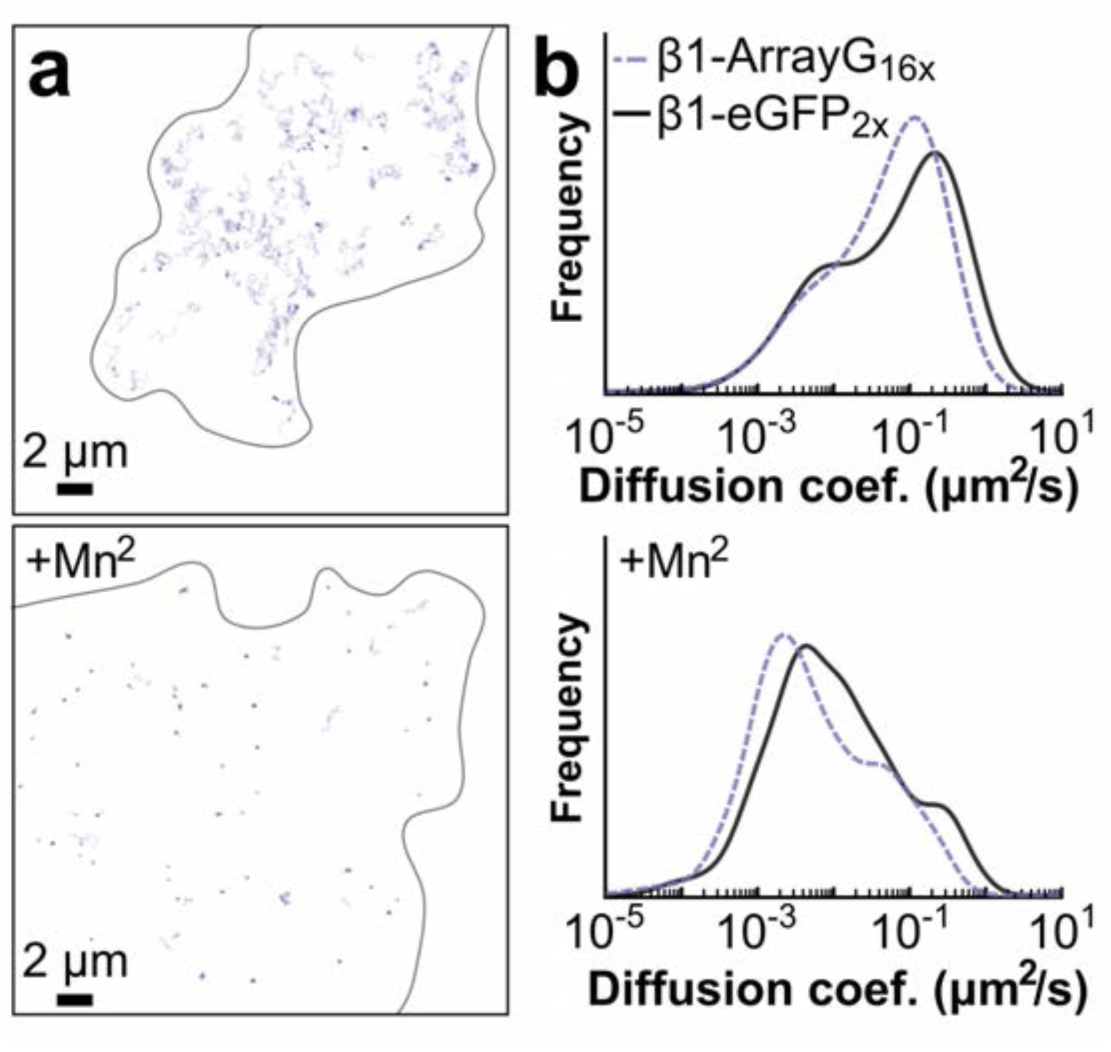
At short time scales, integrin β1-ArrayG_16x_ dynamics match that of β1-eGFP_2x_. (**a**) Trajectories of integrin β1-ArrayG_16x_ in untreated pKOαVβl MEFs (top), and upon 2mM Mn^2+^ treatment (bottom). (**b**) Distribution of diffusion coefficients of β1-eGFP_2x_ (*N* = 4 cells, 659 trajectories) and β1- ArrayG_16x_ (*N* = 9 cells, 2860 trajectory segments) in untreated cells (top), and after 2mM Mn^2+^ treatment ( β1-eGFP_2x_: *N* = 3 cells, 350 trajectories, β1-ArrayG_16x_: *N* = 8 cells, 1388 trajectory segments) (bottom).

We also measured whether β1-ArrayG_16x_ could respond effectively to an external stimulus, by measuring the dynamics of single tagged integrins after addition of Mn^2+^, a potent activator of integrins which enhances global integrin immobilization.^28^ Stimulation with 2 mM Mn^2+^ for 1 hour immobilized eGFP_2x_ and ArrayG_16x_ fusions to a similar extent (trajectories **Figure 5a** and **Supplementary Figure S6**, distributions **Figure 5b**), indicating functionality of the active binding state. Having shown functional complementation of a β1 knockout, correct localization, characteristic diffusion pattern in adhesions, and sensitivity to Mn^2+^ stimulus, we concluded that the ArrayG_16x_ tag did not impair the biological function of integrin β1.

Finally, we compared track length durations of ArrayG_16x_ and eGFP_2x_ fused to integrin β1. The trajectory length distributions between eGFP_2x_ and ArrayG_16x_ revealed a large (~13x) increase in maximum trajectory length (maximum track duration of 8 s vs. 105 s under constant illumination, **Figure 6a**). From 4 cells, we were able to collect 73 tracks each lasting 25 s (500 frames @20 Hz) or longer. We show 6 example of these tracks from a single cell with multiple state switches (as many as 5 transitions) occurring within individual trajectories (**Figure 6b**), compared to the infrequent single transitions seen with the eGFP_2x_ (**Supplementary Figure S6**). We used a simple segmentation analysis to highlight temporary immobilization events, demonstrating the ability to experimentally capture transient sub-trajectory features, which can be used as dynamical signatures of localized function^34^ (**Methods**). We observed both repeated brief immobilization events (**Figure 6b**) and extremely long immobilization events lasting one hundred seconds (**Supplementary Movie S7**). This is consistent with previous estimates of integrin immobilization-lifetimes derived from infrequently sampled sptPALM images (0.25 Hz).^28^

**Figure 6.**
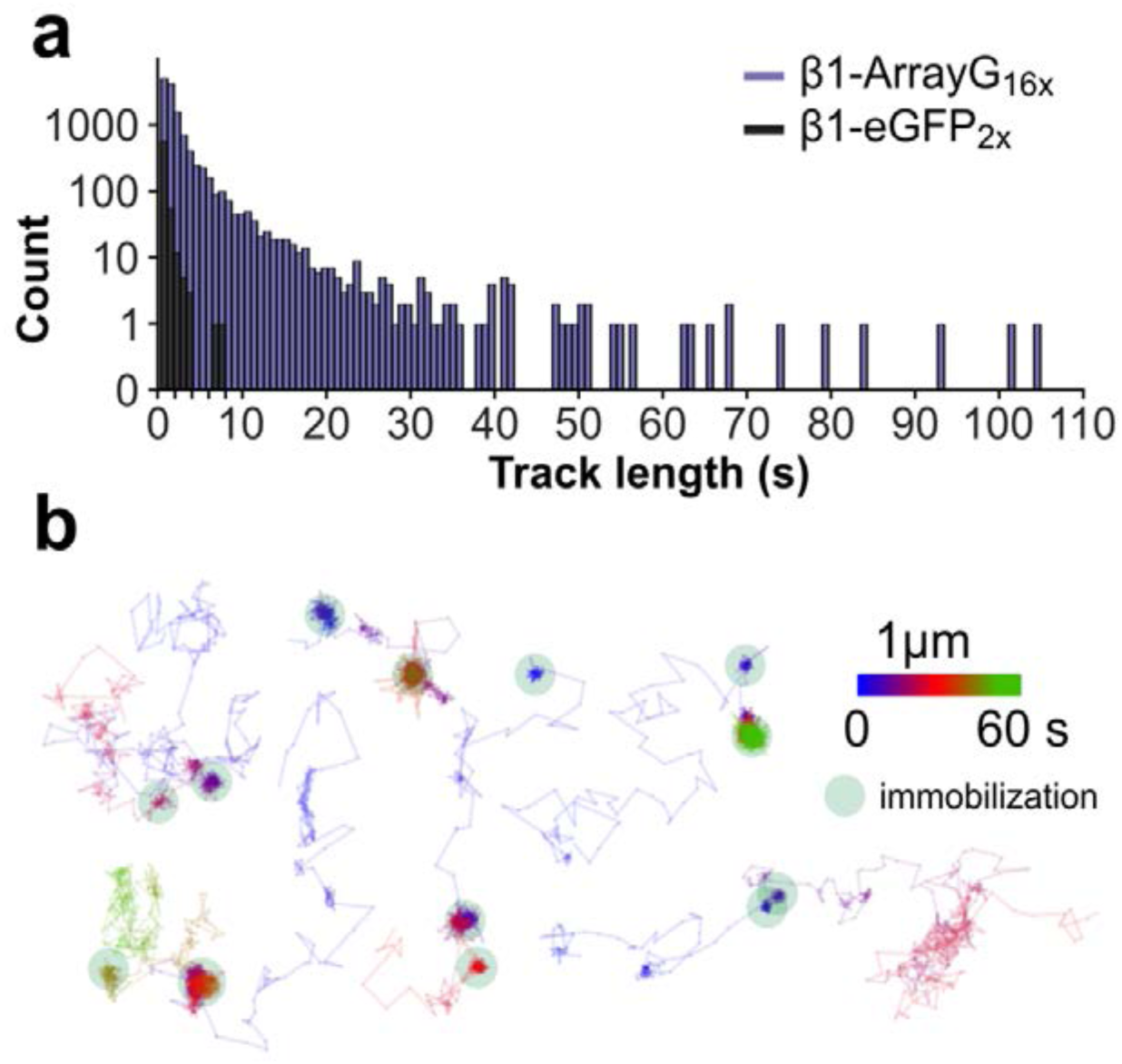
At long time scales, integrin β1-ArrayG_16x_ reveals state-switching dynamics. (**a**) Trajectory length histogram of β1-eGFP_2x_ (659 trajectories from 4 cells) and β1-ArrayG_16x_ (13,423 trajectories from 4 cells). (**b**) Selected β1-ArrayG_16x_ trajectories from a single cell highlighting transitions between immobile and diffusive states. Trajectories are colored by time. Immobilization events are highlighted with a shaded circle.

### NB113 DHFR - a second, orthogonal nanobody array

As noted, our preliminary scanning for functional repeat units revealed di-hydro folate reductase (DHFR) nanobody^35^ as a second potentially suitable protein domain. To generate an array, orthogonal to GBP1, we concatenated repeats of the DHFR nanobody^35^, which binds to *E. coli* DHFR. This second array-binder pair, termed ArrayD, could be used to recruit any fluorescent protein as a fusion to DHFR. We used a simple FRAP assay (**Figure 7a, Supplementary Methods**) to assess the binding efficiencies of ArrayD relative to ArrayG. We found that exchange on ArrayD was ~4x faster than on ArrayG, with recovery half lifetimes of 15 s and 60 s, respectively (**Supplementary Table 2**). Despite a lower reported affinity^35^ and faster exchange rate, ArrayD allowed efficient tracking of single kinesins (**Figure 7b**, **Supplementary Movie S8**). To show that ArrayD_24X_ can be used to generate other colors, we co-expressed KIF560-ArrayD_24X_ with a fusion of mCherry to DHFR (**Supplementary Movie S9**). KIF560-ArrayD_24X_ when co-expressed with mGFP or mCherry gave indistinguishable average speed and track lengths (1.51 μm/s and 1.61 μm/s average speed, 0.94 μm and 1.10 μm average run length, for mGFP-DHFR and mCherry-DHFR, respectively (**Figure 7b**, **Supplementary Figure S7a-d**) Furthermore, by introducing ArrayG_24x_ (with a green binder) and ArrayD_24x_ (with a red binder) in the same cell, we were able to simultaneously track two populations of KIF560 kinesins. These two populations gave indistinguishable average speeds (1.54 μm/s and 1.56 μm/s for mGFP and mCherry, respectively) showing internally consistent results and demonstrated that these arrays are suitable for multicolor imaging (**Figure 7c-d**, **Supplementary Figure S7e, Supplementary Movie S10**).

**Figure 7.**
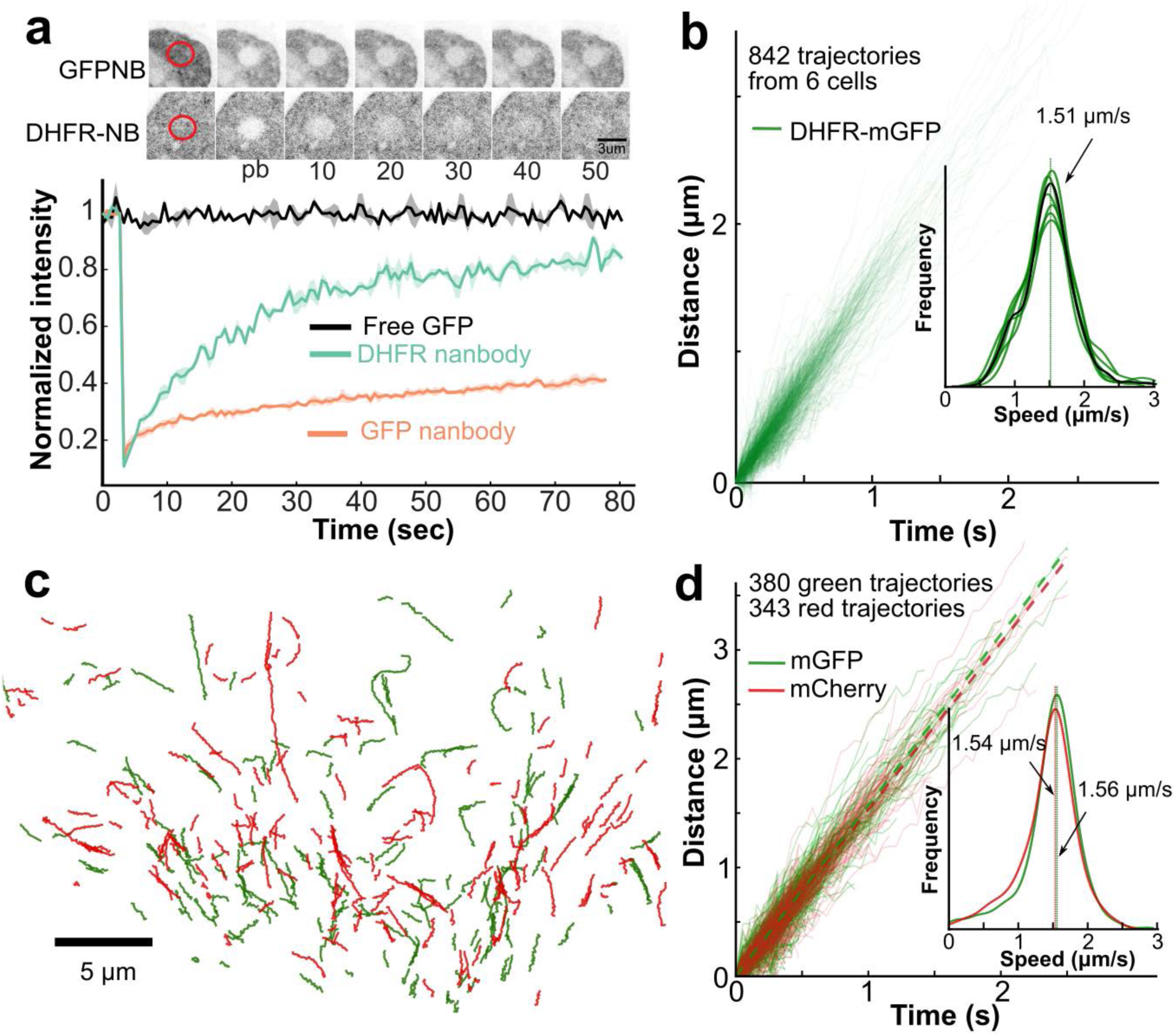
ArrayD, which exhibits faster binder exchange kinetics, provides an alternate prolonged single molecule imaging platform. (**a**) Top: Representative time lapse images of spot bleach and recovery for ArrayG and ArrayD. ~ 10 sec time spacing between images. Bottom: intensity vs. time traces of spot FRAP for different H2B-Array fusions and GFP control (ArrayG: *N* = 3, ArrayD: *N* = 4, Free GFP: *N =* 2). (**b**) Distance versus time for 494 individual trajectories from 7 cells co-expressing KIF560-ArrayD and DHFR-mGFP binder. Inset: distribution of speed. (**c**) Trajectories of kinesin in cells co-expressing KIF560-ArrayG + mwtGFP (green) and KIF560-ArrayD + DHFR-mCherry (red). (**d**) Distance versus time trace of trajectories shown in **c**. Inset: distribution of speed.

## Discussion and Future directions

We have found that protein domains can retain their binding affinity and biochemical activity when concatenated into long arrays. Using such protein domain arrays, we have been able to track single kinesin and integrin β1 molecules inside cells for long durations, revealing complex dynamics of proteins in living systems. Although alternative multi-fluorophore recruitment techniques exist (SunTag^9^, Spaghetti-Monster^36^), only ArrayG has demonstrated the ability to capture high-frequency, long-lasting single-molecule dynamics. It further solves an experimental throughput problem by suppressing background noise through fluorogenicity, thus avoiding forced compartmentalization of binders or cumbersome sorting strategies. Monomeric GFP abrogates aggregation, which we have found to be a potentially major (and yet frequently underemphasized) barrier to precision kinetic and functional single molecule measurements using array-based recruitment technologies. Finally, we have demonstrated prolonged dual color tracking of proteins in living cells by co-expressing two different arrays and corresponding binders. Although illumination times may last as long as 180 seconds before complete photo-bleaching of arrays, the brightness of the ArrayG tag allowed relatively low laser powers to be used, which reduces possible photo-toxicity effects.

In both biological test cases reported here, we found that the target proteins could tolerate array fusions without any detrimental effects on cell viability, cell behavior, or *in cellulo* dynamics of the labeled targets, even when stably expressed in cells. With a typical size of ~1 MDa, ArrayG and D are much larger than most single polypeptides, but are not unusually large compared to cellular multi-protein complexes such as eukaryotic ribosomes (> 3.3 MDa),^37^ cytoplasmic dynein and dynactin ( ~1.2 MDa)^38,39^ and yeast RNA-polymerase II holoenzyme complex (3.4 MDa).^40^ That said, as with any tag, each new ArrayG/D fusion needs to be evaluated for tagging-induced artifacts.

Looking ahead, it might be possible to mix different modular domains within a single protein array allowing for optical or chemical control of array properties. For example, it might be possible to develop arrays with both temporal and spatial control over recruitment of a modifier protein via optically gated binding domains such as LOV^41^ or phytochrome-B (PhyB).^42^ The fluorogenic ArrayG and the high turnover ArrayD may also provide a route for generating refreshable genetically encoded labels with high aggregate photo-stability. GBP1 (enhancer GFP nanobody) derivatives that would allow for faster exchange of mwtGFP at the scaffold would be a first step. This, coupled with ~15 fold brightening of mwtGFP upon binding to GBP1, would allow for sustained fluorescence of the array via exchange with a pool of free dark mwtGFP. GFP nanobodies of varying affinities have already been reported^43^ and could provide a starting point for engineering array tags with fluorogenic potential and relatively lower affinity for mwtGFP, allowing continuous replenishment of fluorescence at the tag. Finally, the fluorogenic ArrayG tag when imaged using appropriate technologies^44^, might allow prolonged single molecule imaging in deep tissues where background fluorescence is particularly inhibitive.

## Materials and Methods

### Array plasmid generation

All mammalian codon optimized Array monomeric unit-DNA were purchased from IDT (integrated DNA technologies) as genes in pUCIDT-Kan plasmid backbone or gBlock gene fragments. All monomeric units had the following structure: (*SalI*)*-6bp spacer-Scaffold unit-flexible Linker-* (*XhoI*)*-6bp spacer-*(*BamHI*). Using the surrounding SalI-XhoI-BamHI combination, we put the monomer unit through several rounds of repeat building, in which we doubled the number of direct repeats with each cloning cycle, as originally shown by Robinett *et al.*^6^ After repeat building, the monomer units were moved into a modified Clontech PEGFP-N1 backbone with a Tet-On 3G Tetracycline inducible promoter in place of the CMV promoter (Tet3G backbone). All repeat-containing plasmids were transformed and maintained in MAX Efficiency Stbl2 competent cells (Thermo Fisher Scientific) grown at 30 °C, as long repeats were unstable in other cell types. For generating double stable cell lines, a version of array backbone was custom built using the PiggyBac Cumate switch system (PBQM812A-1, SBI) where the existing MCS was replaced with a custom MCS, such that Arrays could easily be moved from Tet3G backbone plasmids under the control of cumate promoter. For the binder-FP plasmids, PBQM812A-1 was modified to replace the cumate operator sequence (CuO) with a Tet response element, the EF1-CymR repressor-T2A-Puro cassette was replaced with a custom made EF1-Tet-Activator-T2A-Hygro cassette. Cells efficiently transposed for the array and the binder were puromycin and hygromycin positive respectively and were inducible with cumate (array) doxycycline (binder). Cumate control was leaky enough to produce arrays in amounts that were ideal for single molecule imaging, when cells were uninduced.

Integrin β1 and kinesin source plasmids were purchased from addgene (Addgene numbers 55064 & 15284), vinculin-mCherry was a gift from Johan De Rooij. KIF560 (amino acids 1-560) was generated from full length kinesin. All plasmid sequences are available from NCBI GenBank with the following accession numbers: To be provided by GenBank.

### Cell Culture

HeLa Tet-On^®^ 3G Cell Line (#631183) and U2-OS Tet-On^®^ Cell Line (#631143) were purchased from Clontech. MEF-pKO lines (Shiller *et al.* ^30^) were a gift from Alexander Dunn (Dept. of Chemical Engineering, Stanford University). All mammalian cells were cultured on tissue culture plastic in standard tissue culture incubators held at 37°C with 6.0% CO_2_ using DMEM supplemented with 20% fetal bovine serum and penicillin-streptomycin antibiotics. Before plating pKOαV and pKOαVβ1 mouse embryonic fibroblast cell lines, plastic cultureware was incubated with 5μg/mL fibronectin for at least one hour and then washed with PBS before adding culture media and cells.

### Transfections

All transfections were performed using the Neon Transfection system (Thermo Fisher Scientific). HeLa cells were co-transfected with 0.1-0.2 μg of binder plasmids and 4-8 μg of ArrayG or ArrayD plasmids. We generally used two electroporation pulses (pulse voltage 1,005 V, pulse duration 35 ms). pKO-αVβ1 as well as pKO-αV β1^-/-^ cells were transfected using the following pulse program: 1 pulse, 1,350 V pulse voltage, 30 ms pulse width. Upon transfection, cells were seeded on Mattek dishes coated with fibronectin (Sigma Aldrich) in complete DMEM media without Pen/Strep and with teteracycline free FBS (Thermo Fisher Scientific).

### Cell Line generation

HeLa, U2OS and pKOaV cell lines were generated by transfecting cells with the relevant PiggyBAC plasmids and a plasmid expressing transposase following manufacturer's protocol and then selecting using appropriate antibiotics. For single stable cell line puromycin was the antibiotic of choice where as for double stable cell lines a combination of puromycin and hygromycin was used.

### FACS Analysis

Hela cells stably expressing mwtGFP-P2A-mCherry or mGFP-P2A-mCherry were generated as described above. The cells were analyzed with flow cytometry (BD LSR II) in GFP and mCherry channel along with blank control cells. Only singlets were selected and further analyzed. The ratio of GFP to mCherry per-cell were calculated and displayed after subtracting the background signal from control cells.

### Nuclear compartmentalization assays

The nuclear sequestration assay was developed to test the binding affinity of each binding protein pair in living cells. First, we developed a bulk nuclear compartmentalization “bait-prey” assay (**Supplementary Figure S1**). In this assay, we expressed one partner (the “bait”) from each binder pair as a carboxy-terminal fusion to histone H2B-GFP. Upon expression, these H2B-GFP-bait fusions localized in the nucleus. We co-expressed the bait’s cognate binder (the “prey”) as a fusion to mCherry. All 6 mCherry prey fusions were small enough to passively diffuse into the nucleus.^45^ Images were taken using a Zeiss LSM 700. Image analysis was performed with custom Matlab code. For each cell, a circular ~5 μm ROI was selected in both the nucleus and the cytoplasm, from which the cytoplasm/nucleus (c/n) ratio was calculated. For each condition, 50–150 cells were studied to compute the average and standard deviation. The nuclear/cytoplasmic ratio of mCherry prey provides an estimate of the *in vivo* recruitment potential of the bait. Except for αSyn136–140/Nb2, all binding pairs showed nuclear enrichment (**Sup Mat Table 1, Supplementary Figure S1**). We then constructed long homopolymeric arrays of 24 direct repeats of GBP1 NB, NB113, NBSyn 87, αSyn118–140, EF1, LAM6, and SH3. The domains/epitopes were connected by short glycine/serine linkers and the resulting arrays were fused to H2B. To facilitate nuclear import of these large polypeptides (MW ~1 MDa), a tetrameric repeat of the Importin-β Binding domain (IBB) of Snurportin-α^46^ was inserted between H2B and the array, yielding e.g. H2B-(IBB)_4x_-array fusions. Co-expression of the cognate binder fused to eGFP along with H2B-(IBB)_4x_-array chimeras led to nuclear sequestration of their binder-FP fusions in case of GBP1 and NB113. SH3, LAM6, EF1 and αSyn118–140 failed to recruit binders when concatenated into arrays. This is in no way an exhaustive investigation of the potential of these domains and epitopes, but only a preliminary scan to identify the most promising candidates.

### Single GFP quantification for occupancy estimation method 1: confocal calibration

We performed a multi-step calibration of a Zeiss LSM 700 laser scanning confocal so that we could estimate the number of GFPs within each fluorescent spot. First, we measured the bulk fluorescence of an eGFP solution over a wide variety of microscope settings, varying the PMT gain from 450 to 750 volts, confocal pinhole from 41 to 100 μm, and the laser from 1% to 5% power. We fit all the measurements to an arbitrary three parameter ‘bulk’ model of the form Counts = *A* × laser × (pinhole – *B*) × gain^*C*^, allowing us to estimate the solution’s fluorescence at arbitrary instrument settings (**Supplementary Figure S2**).

We measured the bulk per-molar fluorescence of purified eGFP (CellBioLabs), and 0.04 μm green-fluorescent bead standards (5% w/v FluoSpheres, Molecular Probes) by measuring fluorescence at several fluorophore solution concentrations, and several different microscope settings for redundancy (**Supplementary Figure S2f, g**). To measure bead molarity, which is simply particles per unit volume, beads were immobilized by forming a 5% poly-acrylamide gel within a hemacytometer (Incyto) so they could be counted within a known volume (**Supplementary Figure S2e**). eGFP molarity was measured using a UV-Vis spectrophotometer and the published extinction coefficient at 488 nm.^47^ Using these bulk solution calibrations, we can estimate the per-molar fluorescence of both eGFP and the bead solution at arbitrary gain, laser power, and pinhole settings.

Finally, we measured the average fluorescence intensity from single beads adsorbed to glass. Intensity was extracted from each image as the amplitude of a 2D Gaussian fit to each segmented localization. We measured single bead intensity over a range of instrument settings mirroring those used in the bulk calibration, and fit all the measurements to a similar arbitrary four parameter “single” model of the form Counts = *A* × gain + *B* × laser × (pinhole − *C*) × *D*^gain^ (**Supplementary Figure S2c**). This model allowed us to estimate the fluorescence of a single bead at arbitrary instrument settings.

The ratio of fluorescence from a single emitter to a bulk solution of emitters does not depend on the emitter type, Counts_single_/Counts_bulk_ = *k,* assuming that emitter concentration and imaging conditions are identical.^48^ Thus, we can solve for the fluorescence intensity of a single eGFP as eGFP_single_ = eGFP_bulk_ × (bead_single_/bead_bulk_). The calibrations allow us to estimate single eGFP intensity at experimental gain, laser power, and pinhole settings, using the bulk per-molar fluorescence from both beads and eGFP calculated at the experimental settings from our ‘bulk’ model, and the fluorescence from single beads at the same settings using our ‘single’ model.

### Single GFP quantification for occupancy estimation method 2: HiLo-TIRFM

In a complementary approach to method 1, we estimated the number of eGFPs within single foci using HiLoTIRFM. As HiLo TIRFM is far more sensitive than confocal microscopy, we directly measured the fluorescence from single, purified eGFPs (CellBioLabs) adsorbed to plasma-cleaned glass. We added eGFP to a solution of pH 7.4 PBS while providing simultaneous epi-illumination (0.3 kW/cm^2^) 488 nm laser until we began to see blinking events characteristic of single GFPs non-specifically adsorbing to the surface.^44^ The average eGFP intensity was calculated as the amplitude of a 2D Gaussian fit to each discernable spot. We then imaged ArrayG, without changing any instrument settings, and measured the distribution of single-molecule intensities, again as the amplitude of a 2D Gaussian. We performed a separate eGFP intensity calibration each time each time we quantified ArrayG occupancy.

### TIRF, dual color TIRF and HiLo-TIRFM imaging

Total internal reflection microscopy (TIRF or TIRFM), dual-color TIRFM, and HiLo-TIRFM, were performed on an Olympus CellTIRF system, with independent motor controlled TIRF angles for each fiber-coupled illumination laser, through a 1.49 NA 100x objective and 1.6x Optovar magnifier. Images were recorded on an Andor iXon Plus EMCCD camera at 20 frames per second, unless otherwise noted. Dual-color experiments were imaged through a Photometrics DualView 2 with simultaneous excitation of 488 nm and 561 nm lasers. All dichroics and filters were purchased from Semrock. All experiments were performed within a heated, CO_2_ controlled incubation chamber set to 37 °C and 5%, respectively. Additional temperature control was provided by a collar-type objective heater, also set to 37 °C.

### Single particle tracking

Single particle tracking (SPT) was performed in two steps, all using custom-written Python and C++ code. First, detectable particles in each frame were identified by performing a Laplacian of Gaussian filter and then thresholding the filtered image based on intensity. A 2D Gaussian centered on each identified particle was then refined onto the raw image data using the Levenberg-Marquardt algorithm, with particle x/y position, amplitude, background height, and in some instances standard deviation as free parameters. All particle detection parameters were validated by eye for each dataset, using a custom GUI. The single particle positions in each frame were then linked into single particle time trajectories using a nearest-neighbor type particle tracking heuristic, with a maximum per-frame jump distance parameter. The algorithm iteratively minimized the jump distance between frames, with no assumed momentum. All single particle trajectories were spot-validated by eye. For signal-to-noise ratio calculations, the signal was defined as the peak height of the Gaussian fit above background level, and noise was defined as the standard deviation of a 5 × 5 pixel window centered around the particle location after subtracting fitted Gaussian.

### Kinesin analysis

Starting with raw SPT data, we first removed stuck particles by filtering out tracks that moved less than 0.4 μm/s as measured by the distance between the first and last frame over the total track length. Freely diffusing particles and tracking failures (common at high particle density) were removed by filtering out tracks that turned too sharply, as measured by the maximum track curvature. Using the filtered subset, kinesin speed and run distance were measured by first smoothing each trajectory with a Savitzky-Golay filter to average away particle localization errors, making a one dimensional distance axis. Within each trajectory, each localization was then assigned to the closest interpolated point along the smoothed distance axis, giving run distance as a function of time. Each distance versus time trajectory was fit to a simple linear function to measure run speed, and average speed was measured as the peak of the resultant speed distribution. In high time resolution data, free-diffusing and bound kinesins were distinguished by measuring a sliding window average centered around each time point, +/− 10 frames. Regions with a standard deviation larger than 30 nm were considered diffusing, and lower than 30 nm bound. Speed variation along each bound segment was measured as the first derivative of a Savitsky-Golay filter with a window size of 25 frames, and an order of 2. All analysis was performed using custom code in python.

### Cell spreading assay

To achieve a high expression of the integrin β1-ArrayG constructs, cells were induced with cumate at 5X the working concentration diluted from a 10,000x water-soluble stock solution (SBI 300mg/mL QA150A-1) for at least 3 days before assessing cell spreading. The pKOαV + β1-ArrayG_16x_+mGFP cell line was cultured in the absence of tetracycline using Tet-free FBS, to prevent background induction of the Tet-on inducible control of mGFP. Four hours before performing the spreading assay, mGFP expression was induced with Doxycycline at 2 μg/mL. This was done to promote efficient ER processing and subsequent trafficking of integrin to the membrane without GFP binding, and then subsequently assess functionality of the GFP-occupied array on integrin β1 in the membrane. Under these conditions, cells frequently showed dense assemblies of integrin β1-ArrayG, as shown in Figure 4a (*N =* 3 replicates). Glass bottomed imaging dishes (No. 1 imaging glass, Mattek) were incubated with 10 μg/mL fibronectin in PBS for 2 hours at 37°C. pKOαV cell lines were then seeded on the fibronectin coated dishes and immediately placed on an confocal microscope (Zeiss LSM 700) with a live imaging incubation chamber equilibrated to 37°C and 6% CO_2_. Following a 1-hour equilibration period, cells were imaged using the transmission photomultiplier tube (T-PMT) every 1 hour for 5 hours to obtain bright field images of an 8×7 tile (1145 μm × 1000 μm area). Image analysis took place in three steps: (1) stitching of raw image tiles using on-board Zeiss imaging software, (2) local contrast enhancement of stitched images followed by a Sobbel edge finding filter using ImageJ, (3) Segmentation of cells using custom software written in Python to find cell contours. Each cell contour checked by eye to match the raw images, and manually adjusted if needed. Calculation of mean spreading data used data from cells that remained within the image area and did not divide during the experiment. Data presented represents *N =* 50 to 60 cells per condition (specific numbers given in figure) from two independent replicates. The sample sizes used here are similar to previous measurements.^30^

### MSD analysis

Mean-square displacement (MSD) calculations were done using the *@msdanalyzer* class for MATLAB.^49^ Homogenous drift from the microscope stage was corrected using velocity correlations between individual tracks. After drift correction, the MSD was calculated directly from each trajectory.

### Diffusion coefficient distribution

Integrins exhibit multiple modes of motion that are apparent in the distribution of diffusion coefficients.^28^ We estimated the diffusion coefficient for individual trajectories, rather than ensemble averaged MSD curves, so that we would not loose information about transitions among molecular behaviors. We calculated the distribution of single molecule diffusion coefficients by fitting equation 1 over a specified time scale.

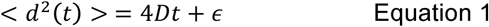

Here, *<d^2^>* is the MSD of an individual trajectory, *t* is the displacement time scale, *D* is the diffusion coefficient and ε is an offset due to localization error. For all datasets, we used *t =* 0.33 seconds to estimate *D.* Images of integrin β1-ArrayG_16x_ were recorded at 20Hz, and integrin β1-eGFP_2x_ images required 30Hz recording to accurately capture its dynamics due to the faster diffusion rate. Because the integrin trajectories from ArrayG-labeled molecules sample long time-scales, it is likely that single molecules switch behaviors over a single trajectory, potentially masking the separate behaviors through averaging. Further, immobilized molecules are more easily tracked, and therefore can have much longer trajectory lengths than mobile molecules. To avoid these biases, we divided long trajectories into a series of non-overlapping 1 second segments, and estimated the diffusion coefficient as described above. Smooth distributions were generated using a normal kernel probability density estimation. To compare the relative diffusion rates of eGFP_2x_ and ArrayG_16x_, the log-scaled diffusion coefficient distribution was fit to a two-Gaussian mixture, giving a rate of 0.25 μm^2^/s for eGFP_2x_ and 0.14 μm^2^/s ArrayG_16x_. Diffusion analysis inside vinculin-mCherry adhesions was performed as follows: (1) registration of GFP and mCherry channels using a rigid-body transformation of the mCherry channel onto the GFP channel from 405 illuminated sample images which produce the same features in each channel, (2) segmentation of time-averaged vinculin-mCherry using a Laplacian of Gaussians filter, and (3) assignment of trajectories that resided within contours of adhesions for 75% of their lifetime as ‘inside’ the adhesion.

### Integrin trajectory segmentation

To accurately detect immobilization events of integrin β1, we employed a previously used detection algorithm in single molecule trajectory analysis.^31,50^ Immobilization events are defined by a spatial component related to localization precision and a temporal-component to ensure the molecule is no longer diffusing. To estimate the localization precision of immobilized integrin β1, we calculated the MSD of trajectories from Mn^2+^ treated cells to ensure tracks are essentially all immobilized. The localization precision, *σ,* is related to the error term of the MSD^51^ (see equation 1): *ϵ* = 4*σ*^2^ We found *σ* = 40 nm (*FWHM* = 2.3σ = 92nm). The probability distribution function of displacements for diffusing particles with a given localization uncertainty is given by:^51^

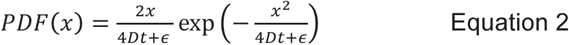

Where *x* is the displacement step, *D* is the diffusion coefficient, *t* is the time step, and ε is the localization uncertainty (4σ^2^). *D* was measured as 0.13 μm^2^/s for integrin β1 ArrayG_16x_ as explained above. We calculated a 99% certainty that a diffusing trajectory would have displaced a distance equal to FWHM after a period of 1 second. We then searched for trajectory segments that remained within a circle of radius FWHM for at least 1 second, and denoted these segments as immobilized.

### Quantifying the dynamics of binder interaction with cognate arrays using FRAP to determine affinity of each array/binder pair

Hela Tet-On 3G cells were co-transfected with H2B-(Snurportin IBB)_4x_-ArrayG or H2B-(Snurportin IBB)_4x_-ArrayD and cognate binder-mGFP plasmids. For line-FRAP experiments, arrays were induced overnight with 1 μg/mL doxycycline. All FRAP analysis was carried out on cells with a typical interphase chromatin architecture, using a Zeiss 700 laser scanning confocal microscope enclosed in an incubation chamber set to 5% CO_2_ and 37 °C. Cells were plated in Mattek glass bottomed dishes coated with fibronectin, and imaged using a 63x oil-immersion objective. Bleaching was carried out using 488 nm laser operating at 100% laser power.

For line-FRAP, we bleached and collected 1-dimensional line scans of 2 pixels wide in 63x oil immersion objective. Average line intensity as a function of time, I(t), was normalized to the average pre-bleach value. The curves were fit with a double exponential function I(t)=A1+A2xEXP(−k1*t)+A3xEXP(− k2*t) (E1) to account for both the fast diffusive free binder-mGFP fraction and the less mobile fraction bound to the cognate scaffold. In case of the control cells expressing mGFP-NLS but no array, the intensity profile was fit with a single exponential function I(t) = A1+A2xEXP(−k1*t) (E2). The fitted parameters were averaged over ~10 repeats to compute means and standard deviations. Half maximum recovery was computed by 1/k in second. For 2D FRAP, we spot bleached a ~2 μm diameter circle, and followed fluorescence recovery. Fluorescence recovery profiles were then analyzed identically to line-FRAP recovery profiles.

### Data and code availability

All datasets and analysis software are available upon request.

## Acknowledgments

This work was partially supported by the National Institutes of Health (NIH) National Institute Of General Medical Sciences (NIGMS)/National Cancer Institute (NCI) Grant GM77856, NCI Physical Sciences Oncology Center Grant U54CA143836, National Science Foundation Graduate Fellowship Program #DGE-114747, and National Institute Of Biomedical Imaging And Bioengineering (NIBIB)/4D Nucleome Roadmap Initiative 1U01EB021237.

## Contributions

R.P.G., W.D., J.M.F, Q.S., and J.T.L. designed research. R.P.G., W.D., J.M.F and Q.S performed research. W.D., J.M.F., Q.S., and R.P.G. analyzed data. R.P.G., W.D., J.M.F, Q.S. and J.T.L. wrote the paper.

